# Immunogenicity of SARS-CoV-2 trimetric spike protein associated to Poly(I:C) plus Alum

**DOI:** 10.1101/2021.10.05.461434

**Authors:** Júlio Souza dos-Santos, Luan Firmino-Cruz, Alessandra Marcia da Fonseca-Martins, Diogo Oliveira-Maciel, Gustavo Guadagini Perez, Victor A. R. Pereira, Carlos H. Dumard, Francisca H. Guedes-da-Silva, Ana C. Vicente Santos, Monique dos Santos Leandro, Jesuino Rafael Machado Ferreira, Kamila Guimarães-Pinto, Luciana Conde, Danielle A. S. Rodrigues, Marcus Vinicius de Mattos Silva, Renata G. F. Alvim, Tulio M. Lima, Federico F. Marsili, Daniel P. B. Abreu, Orlando Ferreira, Ronaldo da Silva Mohana Borges, Amilcar Tanuri, Thiago Moreno L. Souza, Bartira Rossi-Bergamnn, André M. Vale, Jerson Lima Silva, Andrea Cheble de Oliveira, Alessandra D’Almeida Filardy, Andre M. O. Gomes, Herbert Leonel de Matos Guedes

## Abstract

The SARS-CoV-2 pandemic has had a social and economic impact worldwide, and vaccination is an efficient strategy for diminishing those damages. New adjuvant formulations are required for the high vaccine demands, especially adjuvant formulations that induce a Th1 phenotype. Herein we assess a vaccination strategy using a combination of Alum and polyinosinic:polycytidylic acid (Poly(I:C)) adjuvants plus the SARS-CoV-2 spike protein in a prefusion trimeric conformation by an intradermal (ID) route. We found high levels of IgG anti-spike antibodies in the serum by enzyme linked immunosorbent assay (ELISA) and high neutralizing titers against SARS-CoV-2 *in vitro* by neutralization assay, after one or two boosts. By evaluating the production of IgG subtypes, as expected, we found that formulations containing Poly(I:C) induced IgG2a whereas Alum did not. The combination of these two adjuvants induced high levels of both IgG1 and IgG2a. In addition, cellular immune responses of CD4^+^ and CD8^+^ T cells producing interferon-gamma were equivalent, demonstrating that the Alum + Poly(I:C) combination supported a Th1 profile. Based on the high neutralizing titers, we evaluated B cells in the germinal centers, which are specific for receptor-binding domain (RBD) and spike, and observed that more positive B cells were induced upon the Alum + Poly(I:C) combination. Moreover, these B cells produced antibodies against both RBD and non-RBD sites. We also studied the impact of this vaccination preparation (spike protein with Alum + Poly(I:C)) in the lungs of mice challenged with inactivated SARS-CoV-2 virus. We found a production of IgG, but not IgA, and a reduction in neutrophil recruitment in the bronchoalveolar lavage fluid (BALF) of mice, suggesting that our immunization scheme reduced lung inflammation. Altogether, our data suggest that Alum and Poly(I:C) together is a possible adjuvant combination for vaccines against SARS-CoV-2 by the intradermal route.

## Introduction

Intradermal (ID) vaccination has been shown to be a promising strategy to increase the immune response against pandemic viruses such as H5N1 (1), Ebola (2), Vaccinia (3), and SARS-CoV-2 (4). This strategy has already been tested for vaccines against other betacoronaviruses such as SARS-CoV (5, 6), and MERS-CoV (7). Furthermore, the ID route has also been used for clinical trials of MERS-CoV (NCT03721718) and SARS-CoV-2 vaccines (4, 8).

Currently, there are many mRNA and DNA vaccine candidates administered ID against SARS-CoV-2 in mice (9, 10), non-human primates (11) and rabbits (12). Covaxin, an inactivated vaccine developed by Indian pharmaceutical company, Bharat Biotech, was previously tested in mice, rats, and rabbits by the ID route, although the study had focused on the intramuscular (IM) route (13).

The advantages of the ID route include its easy administration through the use of painless microneedles that penetrate the skin, thus increasing vaccination acceptance and coverage, in addition to reducing errors in application and ensuring greater stability of the vaccine formulation (14). Besides, ID immunization has shown success to expand the germinal center (GC) (15), where plasma and memory B cells are regularly generated (16, 17). Potent elicitation of GC response has been demonstrated in experimental mice model of vaccination, which in its turn was highly related to the induction of specific antibodies against SARS-CoV-2, as well as neutralizing antibodies (10,18,19).

Furthermore, ID has already been shown to be a convenient route for the delivery of adjuvants (1). Polyinosinic:polycytidylic acid (Poly(I:C)) is a synthetic analogue of double-stranded (ds) RNA, a molecular pattern associated with viral infections (20). Poly(I:C) triggers a potent type 1 interferon (IFN) response through Toll-like receptor 3 (TLR3) and RIG-I-like receptor (RLR) (21). The activation of these receptors can induce expression of cytokines, chemokines, costimulatory factors, and other dsRNA-dependent systems, resulting in a cellular immune response for viral clearance (20).

A study comparing the role of Poly(I:C) and two derivatives as adjuvants in rhesus macaques demonstrated that treatment with all three Polys were able to induce T cell proliferation (22). Ampligen has been shown to enhance the immunogenicity of an H1N1 influenza vaccine in mice (23). PIKA, a stabilized derivative of Poly(I:C), was reported to be an adequate adjuvant candidate for an inactivated SARS-CoV vaccine, inducing strong anti-SARS-CoV mucosal and systemic humoral immune responses like IgA and IgG (24). Sun et al^25^ reported that Ad5-hACE2-transduced mice, when treated intranasally with Poly(I:C), triggered weight loss control and led to greater viral clearance. Furthermore, Zhao, Wang and Wu showed that HLA-A*0201/Kb transgenic (Tg) mice injected subcutaneously (SC) with Poly(I:C) were able to induce a slight enhancement of SARS-CoV spike peptide-specific CD8^+^ T cells^26^, demonstrating the potential use of Poly(I:C) as a vaccine candidate against SARS-CoV-2.

It was demonstrated that an inactivated vaccine candidate against MERS-CoV using Alum and MF59 adjuvants was able to induce neutralizing antibodies and reduce the viral load in the lungs of experimentally infected mice (22). On the other hand, increased lung immunopathology was observed which was associated with intense cell infiltration of eosinophils (27). However, to reduce eosinophilic infiltration in the lungs of mice, Iwata-Yoshikawa et al^28^ used adjuvant containing Poly(I:C), which also resulted in lower levels of interleukin-4 (IL-4), IL-13, and eotaxin in the lungs. Moreover, Wang et al^29^ demonstrated that the use of Poly(I:C) with other vaccine candidates was able to induce neutralizing and specific antibodies against MERS RBD in mice. Further to this, IFN-γ, IL-4, and IL-2 secreting cells were induced by the Poly(I:C) in comparison to the Alum adjuvant.

Data presented so far suggest that a vaccine given by the ID route using Poly(I:C) and Alum adjuvants could have great potential for a vaccination strategy against SARS-CoV-2. Here, we assess a vaccination strategy using SARS-CoV-2 spike protein in combination with Alum and Poly(I:C) adjuvants by an intradermal (ID) route in a mouse model. After one or two boosts, we found high levels of IgG, IgG1 and IgG2a anti-spike serum antibodies, high neutralizing titers against SARS-CoV-2 and more Spike^+^RBD^+^ B cells in germinal center induced upon the Alum + Poly(I:C) combination. We also found the production of IgG, but not IgA, and a reduction in neutrophil recruitment in the BALF of mice in the challenge with inactivated SARS-CoV-2. Altogether, our data suggest that Alum and Poly(I:C) together is a great strategy for use in combination as adjuvants for vaccines against SARS-CoV-2.

## Material and Methods

### Animals

Female BALB/c mice, 6–8 weeks old (n = 5 per group), were obtained from the breeding facility of UFRJ. All animals were kept in mini-isolators (Alesco, São Paulo, Brazil) and kept under controlled temperature and light/dark cycles of 12 h/12 h, in addition to receiving filtered water and commercial feed (Nuvilab, Curitiba, Paraná, Brazil). The experiments were carried out in accordance with the Ethics Committee on the Use of Animals of the Health Sciences Center of the Federal University of Rio de Janeiro (Comitê de Ética no Uso de Animais do Centro de Ciências da Saúde da Universidade Federal do Rio de Janeiro), under the protocol number: 074/20

### Recombinant SARS-CoV-2 spike glycoprotein used as immunogen

The immunogen used is the whole soluble ectodomain (aminoacids 1-1208) of the spike (S) protein of SARS-CoV-2, containing mutations that stabilize it as a trimer in the prefusion conformation, as first proposed by Wrapp et al^30^. The recombinant HEK293-derived affinity-purified S protein was obtained from the Cell Culture Engineering Laboratory of COPPE/UFRJ, and its purity and antigenicity have already been confirmed in previous works that used it to develop serological tests (31) and equine hyperimmune F(ab’)2 preparations (32). This protein was used in this work to immunize mice and as ELISA antigen to detect anti-SARS-CoV-2 antibodies in samples from immunized animals. Moreover, it was also labeled with Alexa-fluor-467 (similar to APC) and used in germinal center experiments.

### RBD Sequence

RBD (receptor-binding domain): cloning: synthetic gene S1-seq263-685 CoVID-2019_HKU-SZ-005b_2020 (MN975262) (length: 1302 bp) cloned into pET21a(+) using cloning sites NdeI and SalI purchased from Genscript Inc. (USA).

### Expression of recombinant RBD in *E. coli*

Chemically competent E. coli BL21(λDE3) (Novagen, USA) were transformed with 0.2 ng of the pET21a-plasmid, and positive clones were selected in an LB-agar medium containing 100 μg/mL ampicillin at 37 °C overnight. A single positive colony was inoculated in 10 mL of LB medium containing 100 μg/mL ampicillin and this culture was stirred at 220 rpm at 37 °C overnight.

The overnight culture was diluted to 1:100 in 2 L of fresh medium (with antibiotic) and grown at 37 °C until an optical density (O.D._600nm_) between 0.6 to 0.8, when the protein expression was induced with 1 mM IPTG followed by 6 h of expression at 37 °C with 220 rpm stirring. Then, the cells were harvested by centrifugation at 5000 × g for 15 min at 4 °C, and the total-cell lysate was prepared (33).

The pellet was resuspended in 5 mL of buffer A (50 mM sodium phosphate, 200 mM sodium chloride, 1 mM ß-mercaptoethanol, 5 % glycerol, pH 7.0) per gram of pellet, with 1 mM PMSF. Then added 5 mg/mL of lysozyme and stirring for 30 min at 4 °C. After this, 10 µg/mL of DNase A, and 2 mM of MgCl2 were added, and the solution was incubated for 30 min at 4 °C. The total-cell lysate was sonicated using 15 cycles of 15 s on and 45 s off at 300 W (34). The purification of inclusion bodies was performed by centrifugation at 27,000 × g for 60 min at 4°C. Finally, the inclusion bodies were resuspended in buffer B (50 mM sodium phosphate, 200 mM sodium chloride, 1 mM ß-mercaptoethanol, 5% glycerol, 8 M urea, 10 mM imidazole, pH 7.0) overnight at 4 °C^1^. The pellet was resuspended overnight under strong stirring. Next day, the solution was centrifuged at 32,000 x g for 1 h and sequentially filtered through 1, 0.45 and 0.22 µm filters.

### Protein purification

The purification of RBD protein was carried out by affinity chromatography. The protein solubilized in the previous step were subjected to a Ni HP-5 column (5 mL, GE Healthcare, USA), previously equilibrated with 5 CV of Buffer C (buffer C (50 mM sodium phosphate pH 7.0, 0.01 M imidazole and 8 M urea). The nonspecific ligands were removed by washing the column with 5 CV of Buffer C. The elution was performed using a gradient of Buffer C and Buffer D (buffer D, 50 mM sodium phosphate pH 7.0, 0.5 M imidazole and 8 M urea) at a flow rate of 2 mL/min (35). All collected samples were analyzed by SDS-PAGE (36), and the tubes containing the purified interesting protein were pooled and dialyzed in storage buffer.

### Protein labeling with Alexa-fluor 488 and 647

To start labeling proteins with the Alexa kits from ThermoFisher, initially a dialysis was performed to transfer all proteins to PBS buffer. This procedure is recommended by the company to obtain results with a greater labeling efficiency. After this step, the proteins were quantified by Bradford method and then the standard protocol was continued (37). Then, 100 g of protein S and RBD were labeled with kits Alex-fluor-488 (494/519) and Alex-fluor-647 (650/688).

### Cells and virus

African green monkey kidney cells (Vero, subtype E6) were cultured in high glucose DMEM with 10% fetal bovine serum (FBS; HyClone, Logan, UT, USA), 100 U/mL penicillin and 100 μg/mL streptomycin (Pen/Strep; ThermoFisher, MA, USA) at 37 °C in a humidified atmosphere with 5 % CO_2_.

SARS-CoV-2 was prepared in Vero E6 cells. The isolate was originally obtained from a nasopharyngeal swab of a confirmed case in Rio de Janeiro, Brazil (IRB approval, 30650420.4.1001.0008). All procedures related to virus culture were handled in a biosafety level 3 (BSL3) multiuser facility according to WHO guidelines. Virus titers were determined as plaque forming units (PFU)/mL. The virus strain was sequenced to confirm the identity and the complete genome is publicly available (https://www.ncbi.nlm.nih.gov/genbank/SARS-CoV-2/human/BRA/RJ01/2020 or MT710714). The virus stocks were kept in -80°C freezers.

### SARS-CoV-2 inactivation

The SARS-CoV particles were inactivated with β-propiolactone for 24 hours, followed by concentration in 30 % sucrose cushion in 30000 RPM for 1:30 hour. The, the pellet was resuspended in phosphate buffer sodium and storege at - 80 °C.

### Immunization by intradermal route and challenge with inactivated virus

Mice were anesthetized using an induction chamber with an atmosphere saturated with 5% Isoflurane (Cristália^®^, São Paulo, Brazil) in oxygen (Isoflurane Anesthesia Vaporizer System; Harvard Apparatus, MA, USA). Using a 300 µL 30 G syringe (BD Ultra-Fine^TM^), immunization with S Ptn associated with the adjuvants Poly(I:C) HMW (VacciGrade^TM^, 10 mg, batch #VPIC-34-05, InvivoGen, San Diego, CA, USA) and Alum/Alhydrogel 2% (lot #0001657855, InvivoGen, San Diego, CA, USA) was performed in the upper part of the right footpad, in the intradermal region, with visual confirmation of edema after the immunization. The animals received three immunizations, with the same dosage, with an interval of fourteen days between each dose (Supplementary Figure 6). Control animals received only the same volume of PBS. Administrations were as follows: 5 μg of S Ptn per dose (5 μL of a 1 mg/mL solution), 5 μg of Poly(I:C) (1 μL of a 5 mg/mL solution), and 50 % of the final volume was Alhydrogel 2 % (10 μL). PBS was used to make up the final volume to 20 μL. For the challenge, mice received 20 μg of inactivated virus by intranasal route thirteen days after the third dose.

### Intranasal immunization

Mice were immunized three times by intranasal route by installation of 5 µg of Spike protein alone associated with 5 µg Poly (I:C) as adjuvant.

### Antigen-specific antibody responses

Antigen-specific IgG (Cat. No. 1015-05, Southern Biotech, AL, USA), IgA (Cat. No.1040-05, Southern Biotech, AL, USA), IgG1 (Cat. No. 1071-01, Southern Biotech, AL, USA) and IgG2a (Cat. No.1101-01, Southern Biotech, AL, USA) levels were determined by enzyme linked immunosorbent assay (ELISA) using recombinant SARS-CoV-2 S Ptn as the capture antigen. ELISA plates (Corning, MA, USA) were coated with 4 μg/mL of S Ptn in PBS overnight at 4°C. The next morning, the coating solution was discarded and a blocking solution of PBS + 5 % milk (Molico) was added for 1 h. The blocking solution from the ELISA plates was discarded and blood and BALF samples diluted in blocking solution were added for at least 2 h. After this incubation the ELISA plate was washed 5 times with a washing solution consisting of PBS + 0.05 % Tween 20 and then the anti-mouse IgG, IgG1, IgG2a, and IgA-HRP detection antibodies (Southern Biotech, AL, USA) were added for another hour. The plate was washed 7 more times and TMB solution (Invitrogen, MA, USA) was added. The reaction was stopped with 1 N HCl and readed at 450nm. The normalized optical density (O.D.) was made by normalizing data from 4 different experiments using the control groups for that end. The O.D. summatory (Sum) was made by summing the values from normalized O.D. as previously described (25).

### Neutralization assay

To assess the neutralization titer, the serum samples were incubated with 100 PFU of SARS-CoV-2 with serial dilutions of mouse serum for 1 h at 37 °C (to inactivate mouse serum, the samples were heated for 30 min at 56°C). Then, the samples were placed into 96-well plates with monolayers of Vero cells (2 x 10^4^ cells/well) with supernatants for 1 h at 37 °C. Cells were washed and fresh medium with 2 % FBS and 2.4 % carboxymethylcellulose was added. After 72 hours of infection, the monolayer was fixed with formalin 5% and stained with crystal violet dye solution. The cytopathic effect was scored by independent readers. The reader was blind in respect to the source of the supernatant.

### Cell staining for flow cytometry

Cells from lymph nodes (5 x 10^5^) were washed with PBS by centrifugation at 400 *g* for 5 min at 4 °C, blocked with 5 µL/well Human FcX (BioLegend) for 15 min, followed by 5 µL/well of the antibody cocktail and incubation for 30 min at 4 °C in the dark. Cells were washed with a cytometry buffer solution (PBS with 5 % FBS) then fixed with a 4 % formaldehyde (Sigma-Aldrich) for 15 min at 4 °C. Cells were washed, resuspended in cytometry buffer solution, and stored in the dark at 4 °C until acquisition. The following antibodies were used: B220 (anti-B220-PerCP; eBiosciences), GL7 (anti-GL7-PE; BioLegend), CD38 (anti-CD38-PeCy7; BioLegend), RBD-conjugated with Alexa 488, S Ptn-conjugated with Alexa 647, TRCβ (anti-TRCβ-Pacific Blue; BioLegend), CD4 (anti-CD4-FITC; BioLegend), CD8 (anti-CD8-PeCy7; BioLegend), and IFN-γ (anti-IFN-γ-APC; BioLegend). Acquisition of events (100 thousand) was performed on a Becton-Dickinson LSR-II/Fortessa flow cytometer (BD Biosciences, San Diego, CA, USA). The gate strategy was performed based on the selection of cell size (FSC) and composition (SSC). After identifying the main population, a new gate was performed using the FSC-A (area) and FSC-H (weight), where cellular doublets were excluded. The data analyzes were performed using the FlowJo^®^ software vX.0.7.

### Differential BALF cell profile

Immune cells were harvested from the airways via bronchoalveolar lavage (BAL). Briefly, mice were cannulated via a small tracheal incision and the lungs were flushed with 1 mL of sterile PBS. The collected BAL fluids were centrifuged at 1,500 rpm for 5 min at 4°C, and total cells were prepared for flow cytometry staining to determine the number and types of cells. Fc-blocked (1 μg/mL; eBiosciences) BALF cells were stained with anti-mouse SiglecF-PE (0.3 μg/mL; BD Pharmingen), CD11c-APC (0.3 μg/mL; eBiosciences), CD11b-PerCP (0.3 μg/mL; BioLegend), CD19-PECy5 (0.8 μg/mL; eBiosciences), Ly6G-PECy7 (0.8 μg/mL; BioLegend), Ly6C-APC-Cy7 (0.8 μg/mL; BD Pharmingen), and TCRβ-Pacific Blue (0.3 μg/mL; BioLegend). All samples were analyzed on a Becton-Dickinson LSR-II/Fortessa flow cytometer (BD Biosciences, San Diego, CA, USA) and analyzed by using FlowJo software (Tree Star Inc.).

### Data analysis

Results are expressed as mean ± SEM with confidence level of p ≤ 0.05. For multiple comparisons, a one-way ANOVA followed by Tukey multiple comparison test was performed and a two-way ANOVA followed by Bonferroni post-test to compare replicate means by row. Unpaired t-test analysis was done where indicated in the figure legends. Data analysis was performed using GraphPad Prism® 8.0 software.

## Results

### Immunization with Spike protein associated to Alum and/or Poly(I:C) induced high levels of IgG in the serum and in BALF of BALB/c mice

In order to understand whether the vaccine formulations were generating an effective B cell response, we evaluated the different antibody isotypes in the serum and BALF. Mice immunized with S Ptn + Poly(I:C), S Ptn + Alum, or S Ptn + Poly(I:C) + Alum produced higher total specific IgG levels than the group inoculated with S Ptn alone (Figures 1A to 1D). This occurred both at one week after two immunizations (Figures 1A and 1B) and three immunizations (Figures 1C and 1D). Although the group vaccinated with S Ptn presented a higher O.D. Sum than the PBS and Alum + Poly(I:C) groups (Figures 1B and 1D) after the first boost, there were no differences in the O.D. level for each dilution (Figures 1A and 1C). There were very low levels of detectable serum IgG 7 days after prime (Supplementary Figure 1A).

**Figure 1.**
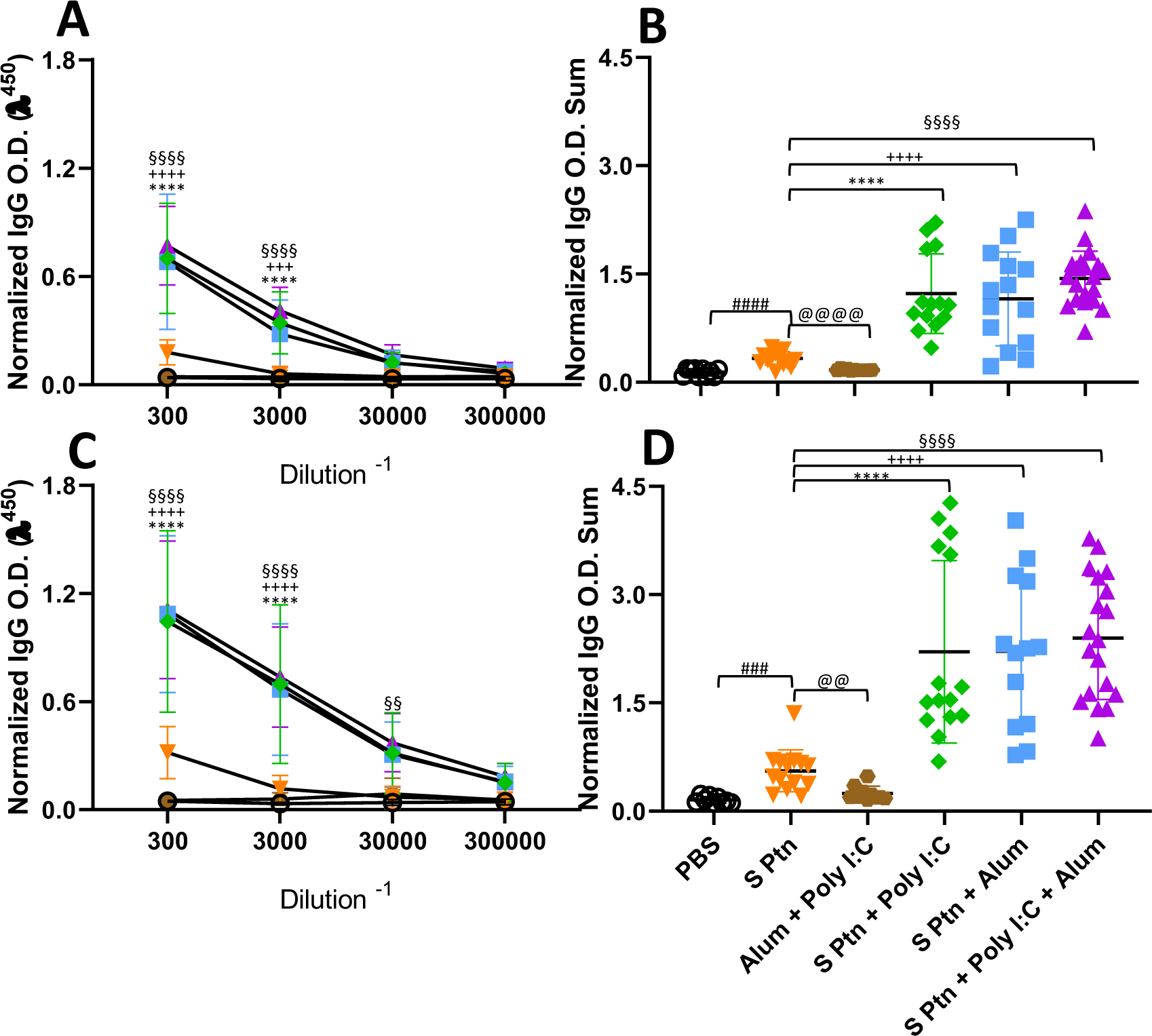
Formulations containing adjuvants induce strong IgG response in the serum. Antigen-specific antibody levels were determined by ELISA and normalized using the control groups. Serum IgG levels were evaluated after one (**A and B**) or two boosts (**C and D)**. The summatory of all dilutions are represented in **B and D**. Data in this figure consists 4 independent experiments normalized using the control groups and shown as mean ± S.D. Groups: Naïve (n=10); PBS (n=15); S Ptn (n=15); Alum + Poly(I:C) (n=11); S Ptn + Poly(I:C) (n=15); S Ptn + Alum (n=15); S Ptn + Poly(I:C) + Alum (n=20). **#** - represents differences between PBS and S Ptn groups; **@** - represents differences between S Ptn and Alum + Poly(I:C) groups; **\*** - represents differences between S Ptn and S Ptn + Poly(I:C) groups; **+** - represents differences between S Ptn and S Ptn + Alum groups; **§** - represents differences between S Ptn and S Ptn + Poly(I:C) + Alum groups; **$**- represents differences between S Ptn + Poly(I:C) and S Ptn + Poly(I:C) + Alum groups; **&** - represents differences between S Ptn + Poly(I:C) and S Ptn + Alum groups; **’**- represents differences between S Ptn + Alum and S Ptn + Poly(I:C) + Alum groups. **A and C** was performed two-way ANOVA followed by Bonferroni post-test and **B and D** was used t-test. *p<0.05, **p<0.01, ***p<0.001, ****p<0.0001.

Since the first interaction between the host and SARS-CoV-2 happens in the lungs, we challenged mice intranasally with inactivated SARS-CoV-2 13 days after the second boost. Twenty-four hours after the viral challenge we euthanized mice and evaluated the antibody levels in the BALF. Our results showed that S Ptn + Poly(I:C), S Ptn + Alum, and S Ptn + Poly(I:C) + Alum formulations induced higher IgG levels in the BALF (Figures 2A and 2B), despite the lack of specific IgA compared to the S Ptn immunization (Figures 2C and 2D). There was no increase in BALF IgA levels between immunized mice versus PBS (FigureS 2C and 2D), indicating no specific IgA response. Furthermore, we observed very low levels of detectable serum IgA even after two and three immunizations in all groups (Supplementary Figures 1B and 1C, respectively).

**Figure 2.**
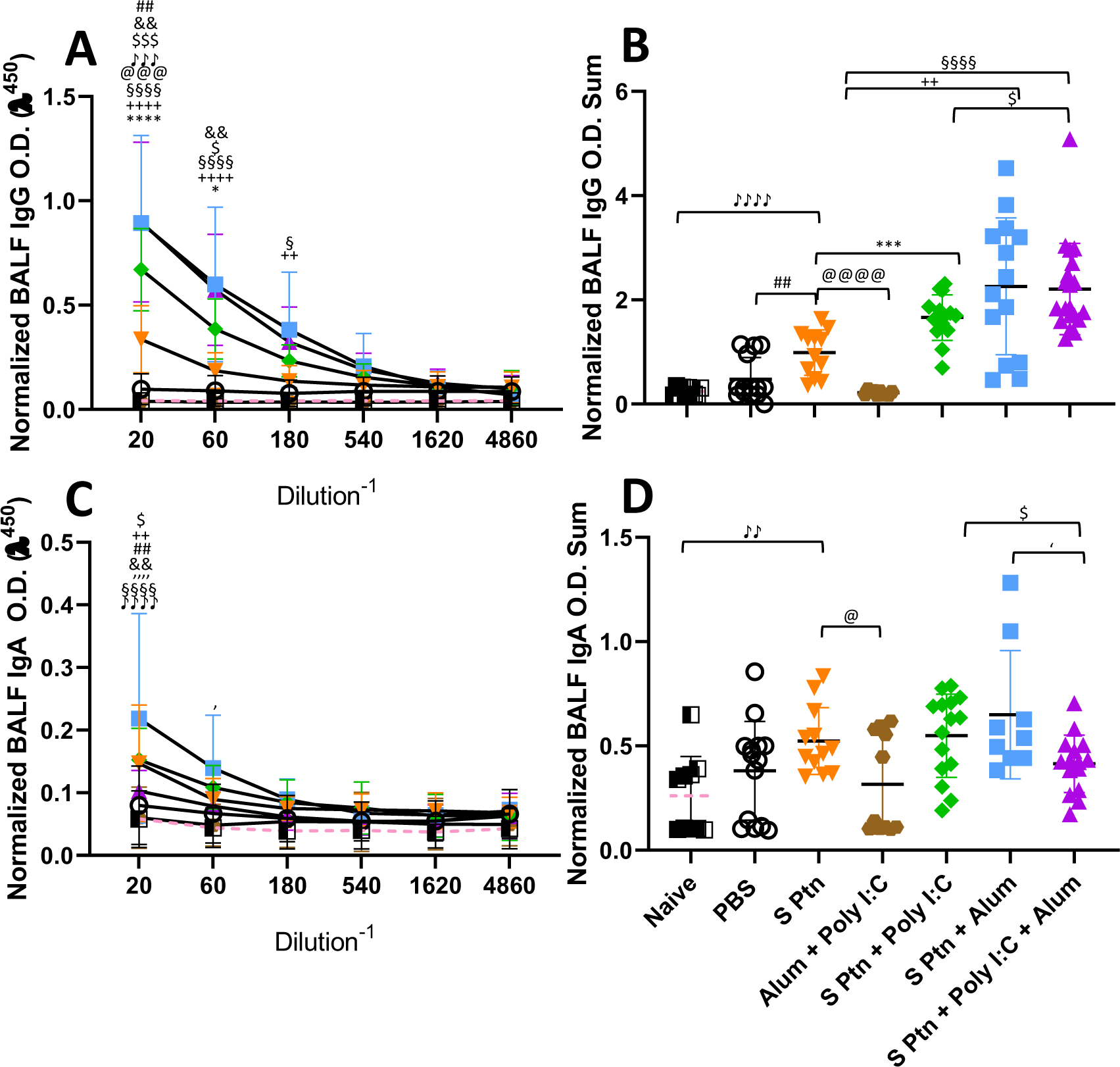
Formulations containing adjuvants induce a strong IgG, but not IgA, response in the BALF. Antigen-specific antibody levels in the BALF were determined by ELISA and normalized using the control groups. Twenty-four hours after the viral challenge we euthanized mice and evaluated in the BALF the IgG (**A and B**) and IgA levels (**C and D**) were determined. The summatory of all dilutions are represented in **B and D**. Data in this figure consists 4 independent experiments normalized using the control groups and shown as mean ± S.D. Groups: Naïve (n=10); PBS (n=15); S Ptn (n=15); Alum + Poly(I:C) (n=11); S Ptn + Poly(I:C) (n=15); S Ptn + Alum (n=15); S Ptn + Poly(I:C) + Alum (n=20). **#** - represents differences between PBS and S Ptn groups; **@** - represents differences between S Ptn and Alum + Poly(I:C) groups; **\*** - represents differences between S Ptn and S Ptn + Poly(I:C) groups; **+** - represents differences between S Ptn and S Ptn + Alum groups; **§** - represents differences between S Ptn and S Ptn + Poly(I:C) + Alum groups; **$** - represents differences between S Ptn + Poly(I:C) and S Ptn + Poly(I:C) + Alum groups; **&** - represents differences between S Ptn + Poly(I:C) and S Ptn + Alum groups; **’**- represents differences between S Ptn + Alum and S Ptn + Poly(I:C) + Alum groups; ♪ - represents differences between Naïve and S Ptn groups. **A and C** was performed two-way ANOVA followed by Bonferroni post-test and **B and D** was used t-test. *p<0.05, **p<0.01, ***p<0.001, ****p<0.0001.

In order to check whether the lack of a specific antibody response in the BALF was due to the immunization route, we performed an intranasal immunization. This route appeared to efficiently increase the amount of IgA (Supplementary Figure 1D) and IgG (Supplementary Figure 1E) in the groups vaccinated with S Ptn + Poly(I:C) in comparison to S Ptn.

### Formulations containing Poly(I:C) induce a higher type 1 response but do not inhibit the type 2 response against SARS-CoV-2 spike protein

The presence of IgG1 is usually related to a type 2 immune response while IgG2a is related to a type 1 response. To understand which response was being induced by the immunizations, we evaluated these IgG subtype (IgG1 and IgG2a) levels in the serum. We found that mice immunized with S Ptn + Poly(I:C), S Ptn + Alum, or S Ptn + Poly(I:C) + Alum produced higher levels of IgG1 compared to mice immunized with S Ptn alone after the first (Figures 3A and 3B) and the second boost (Figures 3C and 3D). Moreover, the S Ptn group presented higher levels of IgG1 than the PBS and Alum + Poly(I:C) groups after the first (Figures 3A and 3B) and second boost (Figures 3C and 3D), but there were no differences in the levels of IgG2a between those groups after either boost (Figures 3E, 3G and 3H). Interestingly, after only one boost, the S Ptn group presented a lower O.D. Sum level than the Alum + Poly(I:C) group (Figure 3F). S Ptn + Poly(I:C) and S Ptn + Poly(I:C) + Alum groups presented higher levels of IgG2a than the S Ptn and S Ptn + Alum groups after one (Figures 3E and 3F) and two boosts (Figures 3G and 3H). Furthermore, mice immunized with S Ptn + Alum had a higher level of O.D. Sum than the S Ptn group after one boost (Figure 3F).

**Figure 3.**
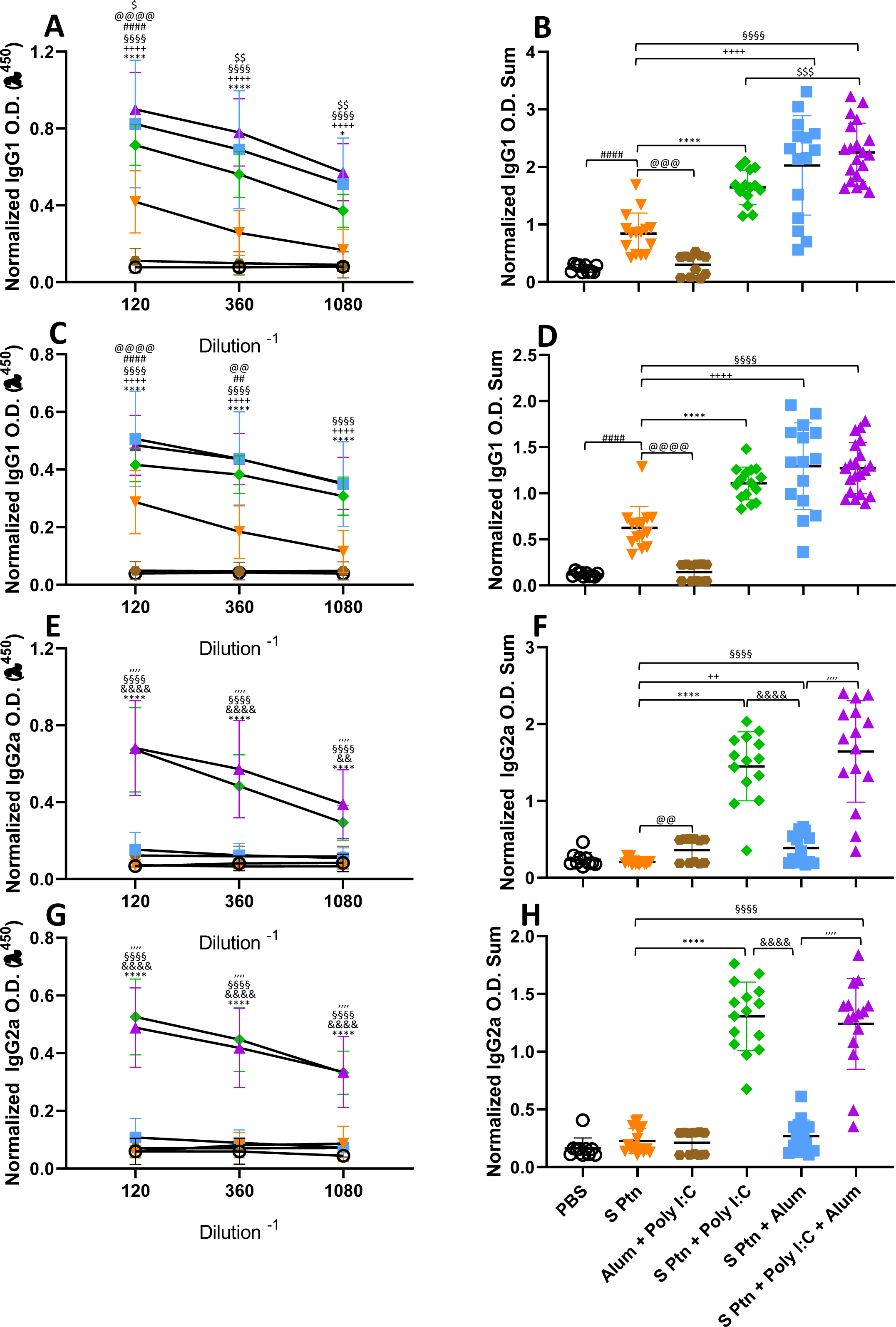
Formulations containing Poly(I:C) were able to induce type 1 response serum antibodies. Antigen-specific antibody levels were determined by ELISA and normalized using the control groups. Serum IgG1 (**A, B, C and D**) and IgG2a (**E, F, G and H**) levels were evaluated after one (**A, B, E and F**) or two boosts (**C, D, G, and H**). The summatory of all dilutions are represented in **B, D, F and H**. Data in this figure consists 4 independent experiments normalized using the control groups and shown as mean ± S.D. Groups: PBS (n=10); S Ptn (n=15); Alum + Poly(I:C) (n=11); S Ptn + Poly(I:C) (n=15); S Ptn + Alum (n=15); S Ptn + Poly(I:C) + Alum (n=20). **#** - represents differences between PBS and S Ptn groups; **@** - represents differences between S Ptn and Alum + Poly(I:C) groups; **\*** - represents differences between S Ptn and S Ptn + Poly(I:C) groups; **+** - represents differences between S Ptn and S Ptn + Alum groups; **§** - represents differences between S Ptn and S Ptn + Poly(I:C) + Alum groups; **$** - represents differences between S Ptn + Poly(I:C) and S Ptn + Poly(I:C) + Alum groups; **&** - represents differences between S Ptn + Poly(I:C) and S Ptn + Alum groups; **’** - represents differences between S Ptn + Alum and S Ptn + Poly(I:C) + Alum groups. **A, C, E and G** was performed two-way ANOVA followed by Bonferroni post-test and **B, D, F and H** was used t-test. *p<0.05, **p<0.01, ***p<0.001, ****p<0.0001.

Together, these data suggest that all adjuvant-containing formulations induce IgG in the serum, but the ones containing Poly(I:C) are able to induce a strong type 1 antibody response against SARS-CoV-2 spike protein, while the ones containing Alum were able to induce slightly higher IgG levels in the BALF (Figure 1E). Immunization with S Ptn alone generated more type 2 antibodies in the serum than the groups containing adjuvants.

### Combination of Spike protein with Alum plus Poly(I:C) induced high neutralization titers

We assessed the *in vitro* neutralizing activity against SARS-CoV-2 in mouse sera collected one week after two and three immunizations (days 21 after first immunization) (Figure 4). We did not observe neutralizing antibodies in the sera of mice immunized with S Ptn, as measured by the neutralizing titers of PRNT_50_ and PRNT_90_. Similarly, naïve and PBS-receiving mice were not able to induce neutralizing antibodies. However, mice immunized with S Ptn associated with the adjuvants Poly(I:C), Alum, or Poly(I:C) + Alum were capable of inducing neutralizing antibodies. The average neutralizing titers of the formulations containing S Ptn + Poly(I:C) + Alum (PRNT_50_ titer of 512 and PRNT_90_ titer of 230.4) were higher than that of S Ptn + Alum (PRNT_50_ titer of 204.8 and PRNT_90_ titer of 96) and S Ptn + Poly(I:C) (PRNT_50_ titer of 108.5 and PRNT_90_ titer of 32). Mice immunized with the adjuvants Poly(I:C) + Alum without S Ptn weren’t able to induce neutralizing antibodies. Moreover, the average of S Ptn + Alum it was superior to the S Ptn + Poly(I:C). These data show that two immunizations are already enough to trigger neutralizing antibodies when S Ptn is combined with the adjuvants Poly(I:C) or Alum, although the mixture of Poly(I:C) + Alum was more efficient.

**Figure 4.**
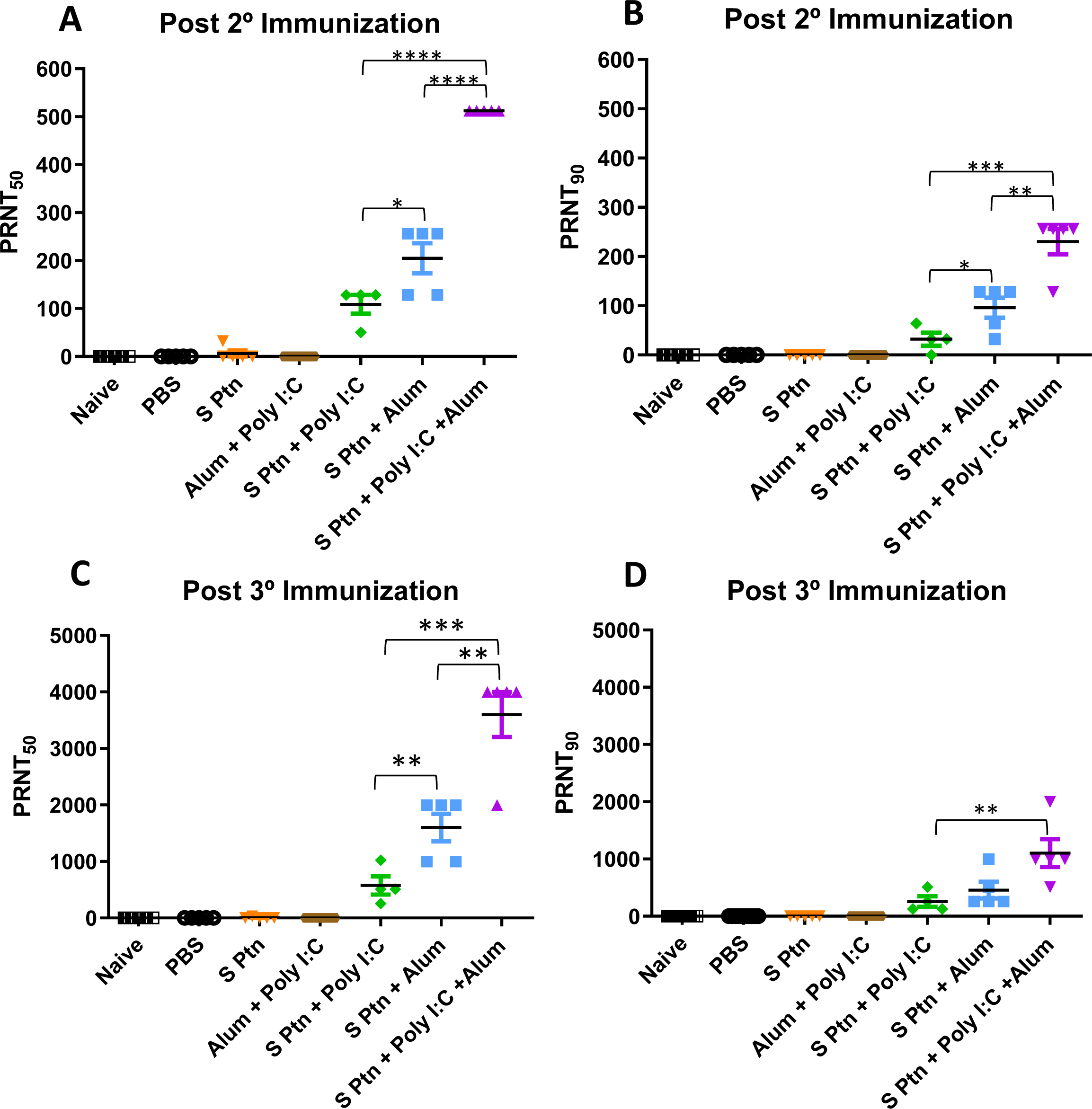
Spike protein associated to Poly(I:C) plus Alum induced high titers of neutralizing antibodies. Titers of neutralizing antibodies were determined *in vitro* by neutralization assay after two and three boosts. PRNT_50_ **(A and C)** and PRNT_90_ **(B and D)** for mice plasma collected 7 days after the second and third immunization. Figure representative of 4 independent experiments and shown as mean ± S.D. with t-test analyses. *p<0.05, **p<0.01, ***p<0.001, ****p<0.0001.

Afterwards, we decided to analyze the neutralizing titers then three immunizations. It was observed that naïve, PBS-receiving mice and mice immunized with the adjuvants Poly(I:C) + Alum without S Ptn weren’t able to induce neutralizing antibodies.

However, mice immunized with S Ptn associated with the adjuvants Poly(I:C), Alum, or Poly(I:C) + Alum were capable of inducing neutralizing antibodies. The average neutralizing titers of the formulations containing S Ptn + Poly(I:C) + Alum (PRNT_50_ titer of 3600) were higher than that of S Ptn + Alum (PRNT_50_ titer of 1600) and S Ptn + Poly(I:C) (PRNT_50_ titer of 576). Moreover, the average of S Ptn + Alum it was superior to the S Ptn + Poly(I:C). Regarding to neutralizing titers with PRNT_90_, the average neutralizing titers of the formulations containing S Ptn + Poly(I:C) + Alum (PRNT_90_ titer of 1102) were higher than that of S Ptn + Poly(I:C) (PRNT_90_ titer of 256).

Taken together, our data show that two immunizations with S Ptn plus Poly(I:C), Alum, or a mixture of Poly(I:C) + Alum is enough to induce neutralizing antibodies, although the mixture presented the highest titers. Moreover, neutralizing titers were higher with three immunizations than those of only two immunizations, therefore having greater potential as a strategy to trigger neutralizing antibodies against SARS-CoV-2.

### Immunization with spike protein associated to Poly(I:C) plus Alum induced high frequencies and numbers of specific B cells in the germinal center

We performed the analysis of lymph node cells draining from the immunization site then the third boost of the groups that received S Ptn alone or S Ptn together with adjuvants. Our results showed that the group that received S Ptn + Poly(I:C) + Alum had an increase in the number of total cells when compared to the group that received only S Ptn (Figure 5A; Supplementary Figure 2). Next, we evaluated the response profile of B cells and observed that there were no differences in the frequency and number of cells between the groups studied (Figures 5B and 5C; Supplementary Figure 3). Within the germinal center, there was an increase both in the percentage and number of cells that were CD38^-^GL7^+^ in the group that received S Ptn + Poly(I:C) + Alum, when compared to the other groups (Figures 5D and 5E). In this cell population, there was an increase in the frequency of RBD^+^S Ptn^+^ cells in the group that received S Ptn + Alum, when compared to the group that received S Ptn + Poly(I:C) and the control groups (Figure 5F). However, we saw that there was a greater number of these cells in the group that received S Ptn + Poly(I:C) + Alum, when compared either to the control groups, the group that received only S Ptn, and the group that received S Ptn + Poly(I:C) (Figure 5G). We also evaluated cells that were RBD^-^S Ptn^+^ and did not observe differences between the groups (Figures 5H and 5I).

**Figure 5.**
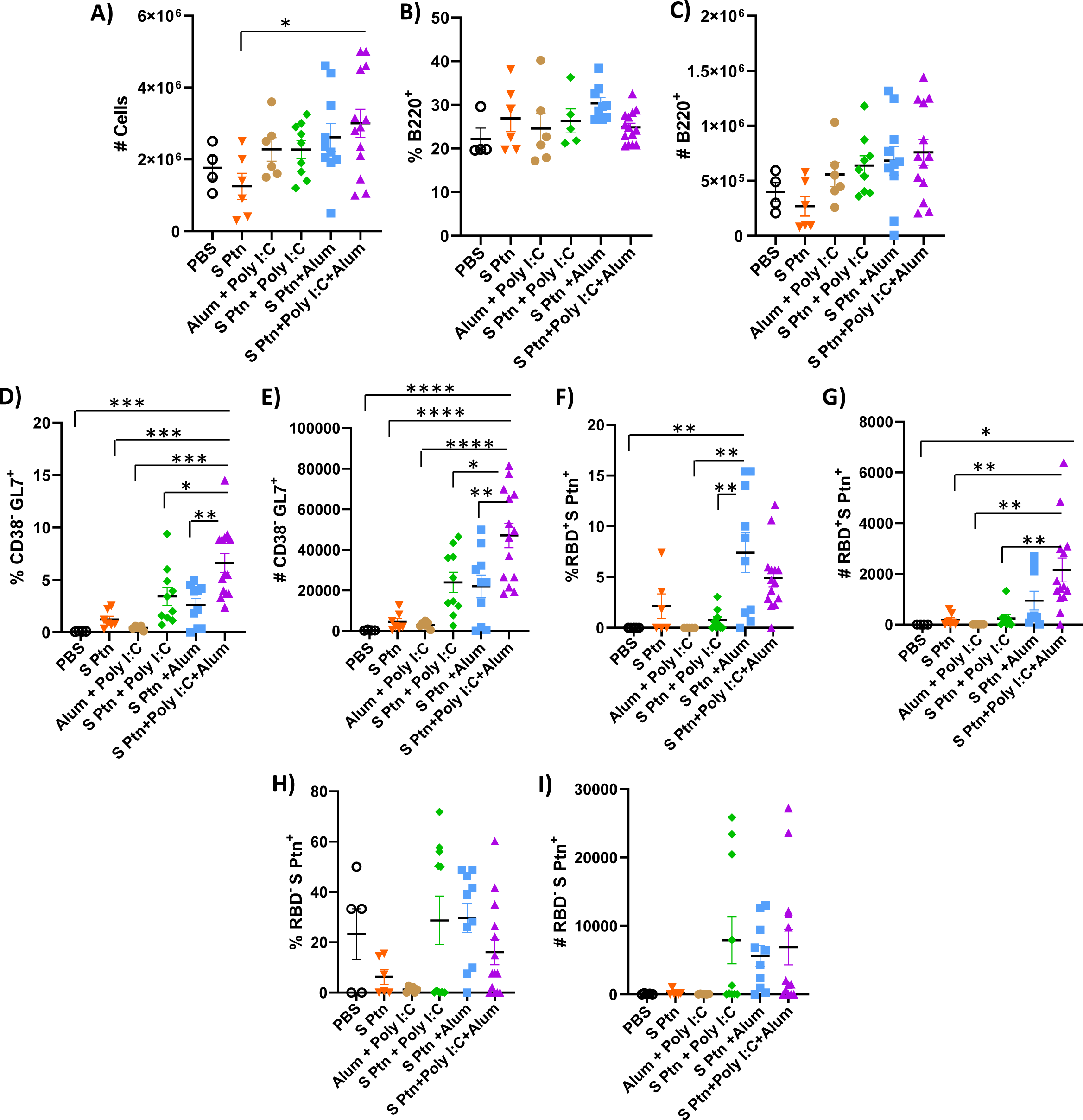
B cell response within the germinal center after immunization. Lymphocytes from the draining popliteal lymph node were analyzed after the third boost following intradermal immunization with S protein alone or with adjuvants (Poly(I:C); Alum; Poly(I:C) + Alum). Controls were performed with PBS or Poly(I:C) + Alum. **(A)** Number of total cells. **(B)** Percentage of B220^+^ cells. **(C)** Number of B220^+^ cells. **(D)** Percentage of CD38^+^GL7^-^ cells. **(E)** Number of CD38^+^GL7^-^ cells. **(F)** Percentage of RBD^+^S Ptn^+^ cells. **(G)** Number of RBD^+^S Ptn^+^ cells. **(H)** Percentage of RBD^-^S Ptn^+^ cells. **(I)** Number of RBD^-^S Ptn^+^ cells. Figure representative of 4 independent experiments and shown as mean ± S.D. and was performed t-test. *p<0.03, **p<0.005, ***p<0.0002, ****p<0.0001. (SEM; n=4-13).

We also continued with the analysis of cells outside the germinal center and noticed that there was a reduction in the frequency of CD38^+^GL7^-^ cells in the group that received S Ptn + Poly(I:C) + Alum in relation to the other groups (Figure 6A). The same was not observed for the number of cells though (Figure 6B). In this cell population, we also observed that the S Ptn + Alum group had an increase in the percentage and number of RBD^+^S Ptn^+^ cells when compared to the other groups (Figures 6C and 6D).

**Figure 6.**
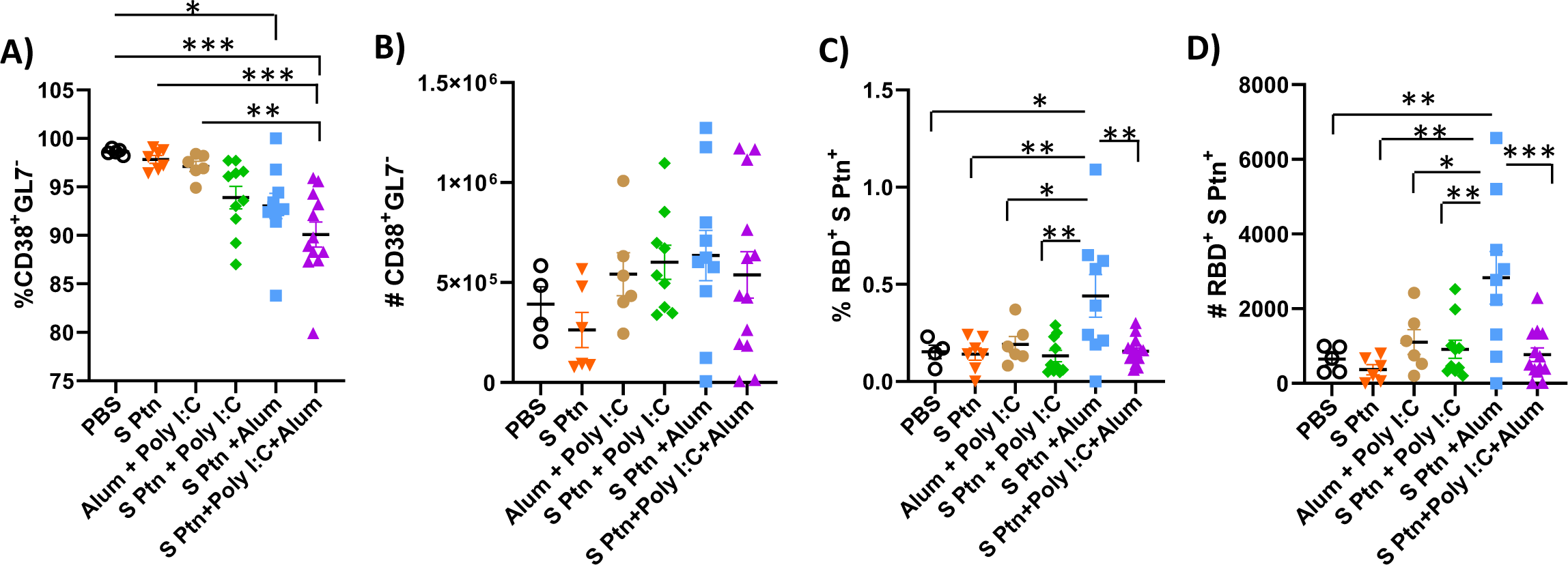
B cell profile outside the germinal center after immunization. After the third boost, lymphocytes from the draining popliteal lymph node macerated after intradermal immunization with S protein alone or with adjuvants (Poly(I:C); Alum; Poly(I:C) + Alum). Controls were performed with PBS or Poly(I:C) + Alum. **(A)** Percentage of CD38^+^GL7^-^ cells. **(B)** Number of CD38^+^GL7^-^ cells. **(C)** Percentage of RBD^+^S Ptn^+^ cells. **(D)** Number of RBD^+^S Ptn^+^ cells. Figure representative of 4 independent experiments and shown as mean ± S.D. and was performed t-test. *p<0.05, **p<0.005, ***p<0.0005. (SEM; n=4-13).

The T cell response was observed for the groups that received S Ptn + Poly(I:C) and S Ptn + Poly(I:C) + Alum as well as the control group performed with PBS. Starting with the CD4^+^ T cells, our data revealed that there was no difference in the frequency of these cells between the groups (Supplementary Figure 4A), but we observed an increase in the number of CD4^+^ T cells in the group that received S Ptn + Poly(I:C) + Alum (Supplementary Figure 4B). We also evaluated the production of IFN-γ by these cells and there were no differences regarding their frequency and number (Supplementary Figures 4C and 4D). Next, we analyzed CD8^+^ T cells and our results showed no differences regarding the frequency of these cells between the groups (Supplementary Figure 4E); however, there was an increase in CD8^+^ T cells in the group that received S Ptn + Poly(I:C) + Alum (Supplementary Figure 4F). The percentage of IFN-y -producing CD8+ cells did not differ (Supplementary Figure 4G); however, we found an increase in these cells in the group that received S Ptn + Poly(I:C) when compared to the control group (Supplementary Figure 4H).

### S protein vaccine formulations induce high neutrophil influx in the BALF of mice after inactivated SARS-CoV-2 challenge

In order to better understand the inflammatory cellular infiltration in the BALF of mice immunized with the different vaccine formulations based on SARS-CoV-2 S Ptn, we performed flow cytometry to distinguish alveolar macrophages (AMs) (SiglecF^+^CD11c^+^), neutrophils (SiglecF^-^CD11b^+^Ly6G^+^), and T cells (SiglecF^-^CD11b^-^TCRβ^+^), 24 hours after challenge with inactivated SARS-CoV-2 (Supplementary Figure 5). We found a pronounced decrease in the percentage (Figure 7A) and absolute numbers (Figure 7B) of AMs in the BALF from non-immunized (PBS) and S Ptn-immunized mice with different combination of adjuvants compared to naïve mice (non-immunized and non-challenged).

**Figure 7.**
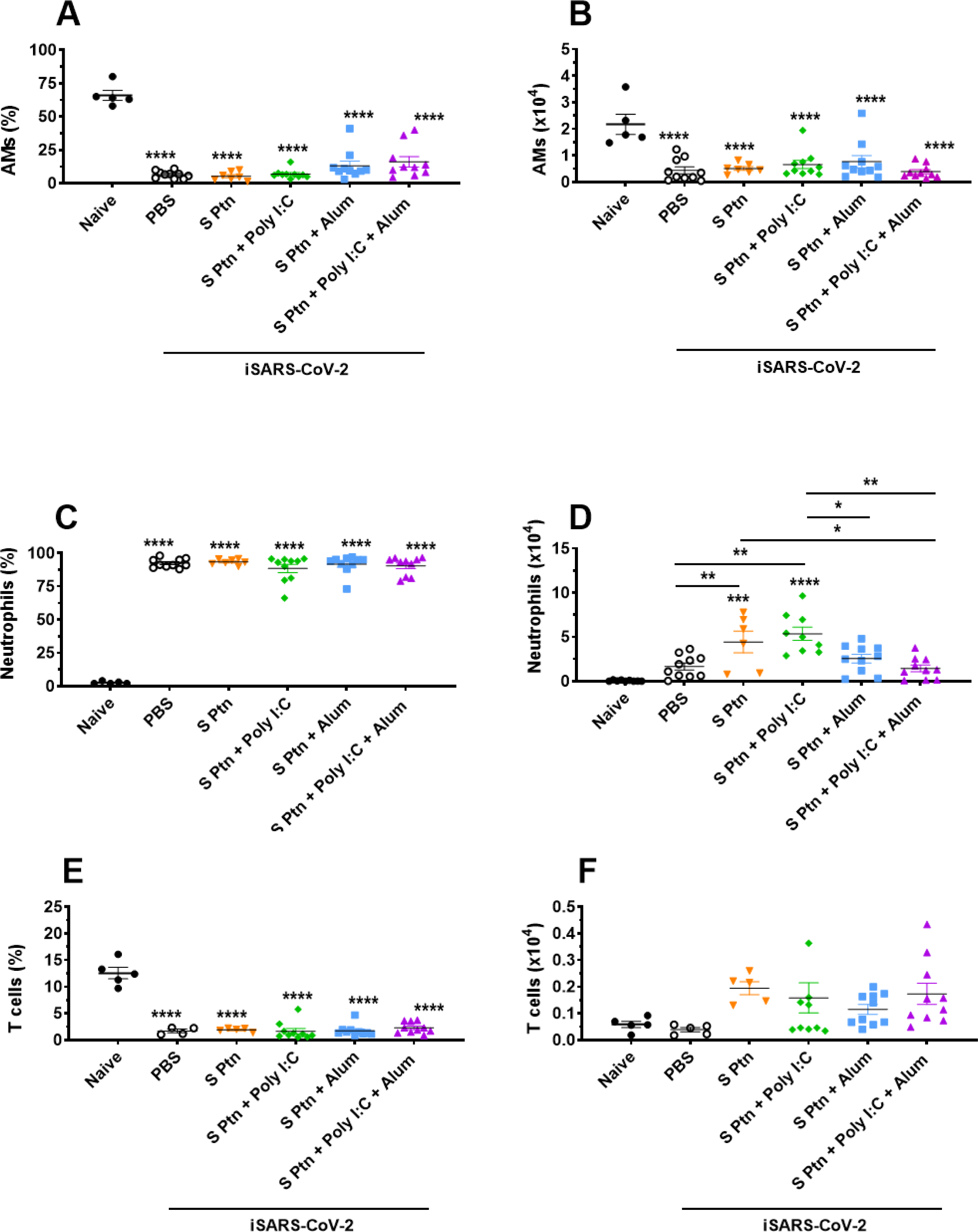
Analysis of BALF immune cell content in mice immunized with S Ptn together with different adjuvants following challenge with inactivated SARS-CoV-2 virus. BALF cells were collected 24 hours after inactivated SARS-CoV-2 (iSARS-CoV-2) virus challenge. The percentage and number of AMs, neutrophils, and T cells from naïve and mice immunized with S Ptn and different adjuvant combinations as gated on flow cytometry plots using markers including SiglecF, CD11c, CD11b, Ly6G, Ly6C, and TCRβ. Percentage of (**A**) AMs (CD11b^+^CD11c^-^ Ly6c^hi^F4/80^+^), (**C**) Neutrophils (CD11b^+^CD11c^-^Ly6c^+^F4/80^-^), and (**E**) T cells (CD11b^+^CD11c^-^Ly6c^+^F4/80^-^) in the BALFs. Absolute numbers of (**B**) AMs, (**D**) Neutrophils, and (**F**) T cells in the BALFs. Data are presented as the mean ± SEM of three pooled experiments and analyzed by one-way ANOVA with Tukey’s post hoc test, *p<0.05, **p<0.01, ***p<0.001, ****p<0.0001.

COVID-19 severity is marked by neutrophil and T cell imbalance in blood samples of patients (38, 39). In addition, elevated numbers of neutrophils are observed in the nasopharyngeal epithelium and later in the more distal parts of the lungs upon SARS-CoV-2 infection (40, 41). In our model, we found a higher percentage of neutrophils (Figure 7C) and lower percentage of T cells (Figure 7E) in the BALF of mice immunized with S Ptn together with the different combinations of adjuvants compared to naïve mice. Interestingly, we also observed that injection itself with either PBS or S Ptn alone increased the percentage of neutrophils and decreased the percentage of T cells in the BALF of challenged mice compared to the percentages found in naïve mice (Figures 7C and 7E). On the other hand, the absolute numbers of neutrophils were quite variable among non-immunized and S Ptn-immunized mice. Specifically, we found that S Ptn alone and S Ptn + Poly(I:C) immunization induced an enhancement in the numbers of neutrophils when compared to non-immunized BALF of SARS-CoV-2 challenged mice. In addition, S Ptn without adjuvant increased the numbers of neutrophils when compared to the S Ptn + Poly(I:C) + Alum group. Finally, S Ptn + Poly(I:C) induced an enhancement in the numbers of neutrophils when compared to the S Ptn + Alum and S Ptn + Poly(I:C) + Alum groups (Figure 7D). No statistically significant increases were found for the absolute numbers of T cells among groups of non-immunized and S Ptn-immunized mice with different combinations of adjuvants. However, we found a tendency of increasing numbers of T cells in mice immunized with S Ptn together with different combinations of adjuvants compared to naïve and non-immunized mice (PBS) (Figure 7F).

## Discussion

The ID route has recently been shown to be an optimal immunization strategy against SARS-CoV-2 due to its ability to stimulate Antigen Presenting Cells (APCs), such as Dendritic cells (DCs) and macrophages, in the dermis, which presents high vascularization, facilitating the migration of cells to secondary lymphoid organs for T and B cell activation (42). One of the main advantages of the ID route is that, due to the presence of a large number of APCs, it allows the administration of a lower dose of antigens and adjuvants is required to generate immune responses, unlike those necessary for the IM and SC routes (43-45).

To this day, there have been two vaccines approved for use in humans against SARS-CoV-2 that use the adjuvant Alum in the formulation together with purified virus inactivated by β-propiolactone, PiCoVacc (46) and BBIBP-CorV (47). Zhang et al^46^ observed that various doses of PiCoVacc mixed with Alum (0, 1.5, 3, or 6 μg per dose) in BALB/c mice induced neutralizing antibodies against S Ptn after two immunizations by the IM route (7, 14, 21, 28, and 35 days). The animals immunized with Alum only did not induce neutralizing antibodies. In the experimental trials of BBIBP-CorV, Wang et al^48^ evaluated the immunization of 0.5 mL of vaccine (2, 4, or 8 μg per total dose) containing Alum (0.45 mg/mL) following the protocol with one-dose (D0), two-dose (D0/D21), and three-dose (D0/D7/D14) in BALB/c mice administered intraperitoneally. They showed that 28 days after the first immunization the three-dose immunization program led to higher levels of neutralizing antibodies than both the one- and two-dose protocols. In addition, they demonstrated that the two-dose immunization protocol was also able to induce neutralizing antibodies at 7 days after the second immunization (day 21). Our data demonstrated similar results, showing that the second immunization via the ID route with S Ptn associated to Alum, Poly(I:C), and Alum plus Poly(I:C) is already sufficient to induce neutralizing antibodies and that a third immunization is capable of inducing even more mainly for the combination with Alum plus Poly(I:C).

Although Alum is acknowledged for its ability to induce a Th2 response (49), many studies regarding SARS-CoV-2 vaccine development have reported a Th1 response directed by Alum (50-52), which was associated to the TLR9 adjuvant CpG + Alum coupled activity in some cases (53-55). Poly(I:C) is a TLR3 agonist and formulations of mixed adjuvant containing Alum + Poly(I:C) have been shown to elicit a T cell immune response. This response was observed by the increase of antigen-specific IgG1 and IgG2a levels, related as well to maturation and activation of dendritic cells. The same study observed a similar response for a formulation of Alum + CpG (56). Chuai et al^57^ also demonstrated that immunization of mice with Alum + Poly(I:C) formulation in conjunction with the S protein from Hepatitis B virus induced increased levels of IgG, as well as IFN-γ, and IL-2, which are related to a Th1 immune response. Along with that, researchers have studied a derivative of Poly(I:C), poly-ICLC (Hiltonol), which is already being tested in clinical trials (NCT04672291), and has shown to be efficient in protecting BALB/c mice in a lethal SARS-CoV infection model (58). Moreover, Smith et al^59^ demonstrated that immunization of BALB/c mice with synthesized vaccine peptides was capable of inducing IFN-γ release in response to predicted T cell epitopes in mice vaccinated with peptides + Poly(I:C) rather than Poly(I:C) alone.

We therefore suggest our formulation of S Ptn + Alum + Poly(I:C) is capable of stimulating Pattern Recognition Receptors, leading to activation of the innate immune response and an antiviral response, such as induced IFN-γ production and increased T cell activation. An adaptive immune response was also observed, with high production of antigen-specific IgG, as well as higher levels of GC B cells specific for both RBD and S Ptn. Our formulation of S Ptn + Alum + Poly(I:C) was capable of driving a sustained type 1 immune response, with the production IFN-γ and IgG2a, as well as inducing higher levels of type 2 immune response antibodies like IgG1 compared to the group immunized with S Ptn alone.

Nashini et al^60^ analyzed an immunization strategy against SARS containing Alum mixed with other adjuvants by the IM route in BALB/c mice. They noted that the formulation containing Alum (100 μg) + Poly(I:C) (50 μg) was able to induce more neutralizing antibodies than a formulation containing either only Poly(I:C) or no adjuvants at 28 days after immunization in old and young mice. This response was still observed at 210 days post-immunization with the formulation containing Alum + Poly(I:C). The use of Alum (50 μg) and Poly(I:C) (50 μg) has also been proven to induce neutralizing antibodies against MERS-CoV (29).

It was demonstrated by Lederer K, et al^61^ that after mRNA immunization in humans induced B cells that recognize Spike^+^ RBD^+^ (at the same time) and Spike^+^ RBD^-^ in the germinal centers of the draining lymph nodes. We also observed after immunization using spike proteins associated to adjuvants Alum, or Poly(I:C) or the combination with alum:Poly(I:C) high levels of B cells in GC that recognize Spike^+^ RBD^-^ and double positives Spike^+^ RBD^+^ were found. We also observed that B cells outside GC can recognize Spike^+^ RBD^+^ and Spike^+^ RBD^-^, however, with fewer recognition in comparison to cells from GC. Taken together, these data may lead to the implication that reactions on the GC are highly required for the formation of neutralizing antibodies. However, more studies to evaluate the importance of somatic hypermutation to generate high-affinity GC B cell clones are required to confirm the role of GC to formation of neutralizing antibodies.

Understanding the efficacy of a vaccine also requires a comprehensive assessment of cellular events that occur after its administration. The literature on COVID-19 vaccine efficacy is focused on the important parameters of antibody production and T cell activation. However, one of the hallmarks of COVID-19 severity is the elevated numbers of neutrophils in the blood samples (38, 39) as well as in the nasopharyngeal epithelium and later in the more distal parts of the lungs upon SARS-CoV-2 infection (38, 39). In addition, a recent study reinforced the importance of reducing neutrophil recruitment to the lungs to prevent severe forms of COVID-19 (62). Here, we found a higher recruitment of neutrophils to the lungs of S Ptn-immunized mice following SARS-CoV-2 challenge. However, when S Ptn was combined with Poly(I:C) + Alum adjuvants, we found lower numbers of neutrophils in the BAL of these immunized mice compared to all the adjuvants combination tested. Based on the reduction of neutrophils we wonder the possibility of the impact of vaccination using the combination of S Ptn + Poly(I:C) + Alum could reduce inflammatory response caused by neutrophils. Futures experiments using challenge can confirm this hypothesis.

Taking these data together, we suggest our formulation of S Ptn + Alum + Poly(I:C) is capable of inducing neutralizing antibodies against SARS-CoV-2 at rates higher than formulations with single adjuvant or no adjuvants at 21 days and 35 days after the second and third immunizations, respectively. The combination of both Alum and Poly(I:C) adjuvants together with S Ptn as antigen are good candidates for a vaccine against COVID-19.

## Ethics Statement

The animal study was reviewed and approved by the Animal Ethical Committee of Federal University of Rio de Janeiro (number 074/20).

## Author Contributions

HLMG, ACO, AMO and JLS designed the study. JSS, LFC, AMFM, DOM, GGP, VARP, ADF and HLMG wrote the manuscript. JSS, LFC, AMFM, DOM, GGP, VARP and CHR performed the experiments. Assistance in experiments: FHGS, ACVS, MSL, JRMF, KGP and BRB. JSS, LFC, AMFM and ADF analyzed the data. JSS and DOM – Immunization and Challenge. JSS, LFC and AMFM - Flow cytometry. LFC - ELISA. ADF, MSL, JRMF and KGF - BALF. LC, DASR, MVMS, OF, RSMB and AMO - S Pnt labeling, RBD. RGA, TML, FFM and DPA - S Ptn production. Final review of the text by HLMG, JLS, ACO, AMO and AMV. Scientific discussion performed by JSS, LFC, AMFM, DOM, GGP, JLS, ACO, AMO, AMV, BRB and HLMG. All authors contributed to the article and approved the submitted version.

## Acknowledgment

We thank Professor Leda Castilho from COPPE/UFRJ to donate Spike protein for this study. We also thank Serrapilheira, FAPERJ and CNPq for financial support.

---

**Supplementary Figure 1 – The prime was not a good inducer of IgG, and IgA levels were very low after boosts.** Antigen-specific antibody levels were determined by ELISA and normalized using the control groups. Serum IgG levels (**A**) was evaluated 1 week after prime. Serum IgA levels (**B and C**) were evaluated after one (**B**) or two boosts (**C**). BALF IgA (**D**) and IgG (**E**) levels. Data in **A, B and C** are representative of 4 independent experiments and are shown as mean ± S.D. Data in **D and E** are from a single experiment and are shown as mean ±S.D. Groups: PBS (n=5); S Ptn (n=5); Alum + Poly(I:C) (n=6); S Ptn + Poly(I:C) (n=5); S Ptn + Alum (n=5); S Ptn + Poly(I:C) + Alum (n=5). **\*** - represents differences between S Ptn + Poly(I:C) (intradermal) and S Ptn + Poly(I:C) (intranasal) groups;

**Supplementary Figure 2 - Gating strategy of B cells in draining lymph nodes.** Lymphocytes from the draining popliteal lymph node were analyzed after the third boost as follows: (A) Cells FSC x SSC. (B) Single cells - FSC-A x FSC-H. (C) B220^+^ (PerCP x FSC-A). (D) CD38^-^GL7^+^ (PE-Cy7 x PE). (E-F) RBD^+^S Ptn^+^ (FITC x APC). (G) FMO GL7. (H) FMO RBD. (I) FMO S Ptn.

**Supplementary Figure 3 - Dot plot of B220^+^ cells.** Lymphocytes isolated from the draining popliteal lymph node after the third boost by intradermal immunization with S Ptn alone or with adjuvants (Poly(I:C); Alum; Poly(I:C) + Alum). Controls were performed with PBS or Poly(I:C) + Alum. Dot plot of B220^+^ cells (PerCP x FSC-A).

**Supplementary Figure 4 - Profile of CD4^+^ T and CD8^+^ T cell response after immunization.** Lymphocytes isolated from the draining popliteal lymph node after the third boost by intradermal immunization with S Ptn plus Poly(I:C) or Poly(I:C) + Alum. Controls were performed with PBS. (A) Percentage of CD4^+^ T cells. (B) Number of CD4^+^ T cells. (C) Percentage of IFN-γ^+^CD4^+^ T cells. (D) Number of IFN-γ^+^CD4^+^ T cells. (E) Percentage of CD8^+^ T cells. (B) Number of CD8^+^ T cells. (C) Percentage of IFN-γ^+^CD8^+^ T cells. (D) Number of IFN-γ^+^CD8^+^ T cells. *p<0.05 (T Test). (SEM; n=5-15).

**Supplementary Figure 5 -** Gating strategy for the flow cytometric detection of macrophages, eosinophils, T cells, and neutrophils in BALF. BALF cells were first gated on a forward scatter/side scatter (FSC-A/SSC-A) and then on single cells using FSC-A/FSC-H. A sequential gating strategy was used to identify cellular populations expressing specific markers: alveolar macrophages (AMs) (SiglecF^+^CD11c^+^), eosinophils (SiglecF^+^CD11c^-^), T cells (CD11b^-^TCRβ^+^), and neutrophils (CD11b^+^Ly6C^-^Ly6G^+^).

**Supplementary Figure 6 – Protocol of Immunization.** The animals received three immunizations, with the same dosage, with an interval of fourteen days between each dose. Blood samples was collected after one and two boosts. Then, mice were challenge with inactivated SARS-CoV-2 and 24h later the mice were euthanized.

