## Supplementary material for "Immunogenicity of SARS-CoV-2 trimetric spike protein associated to Poly(I:C) plus Alum": Supplemnetary figures

### Supplementary Figure 1

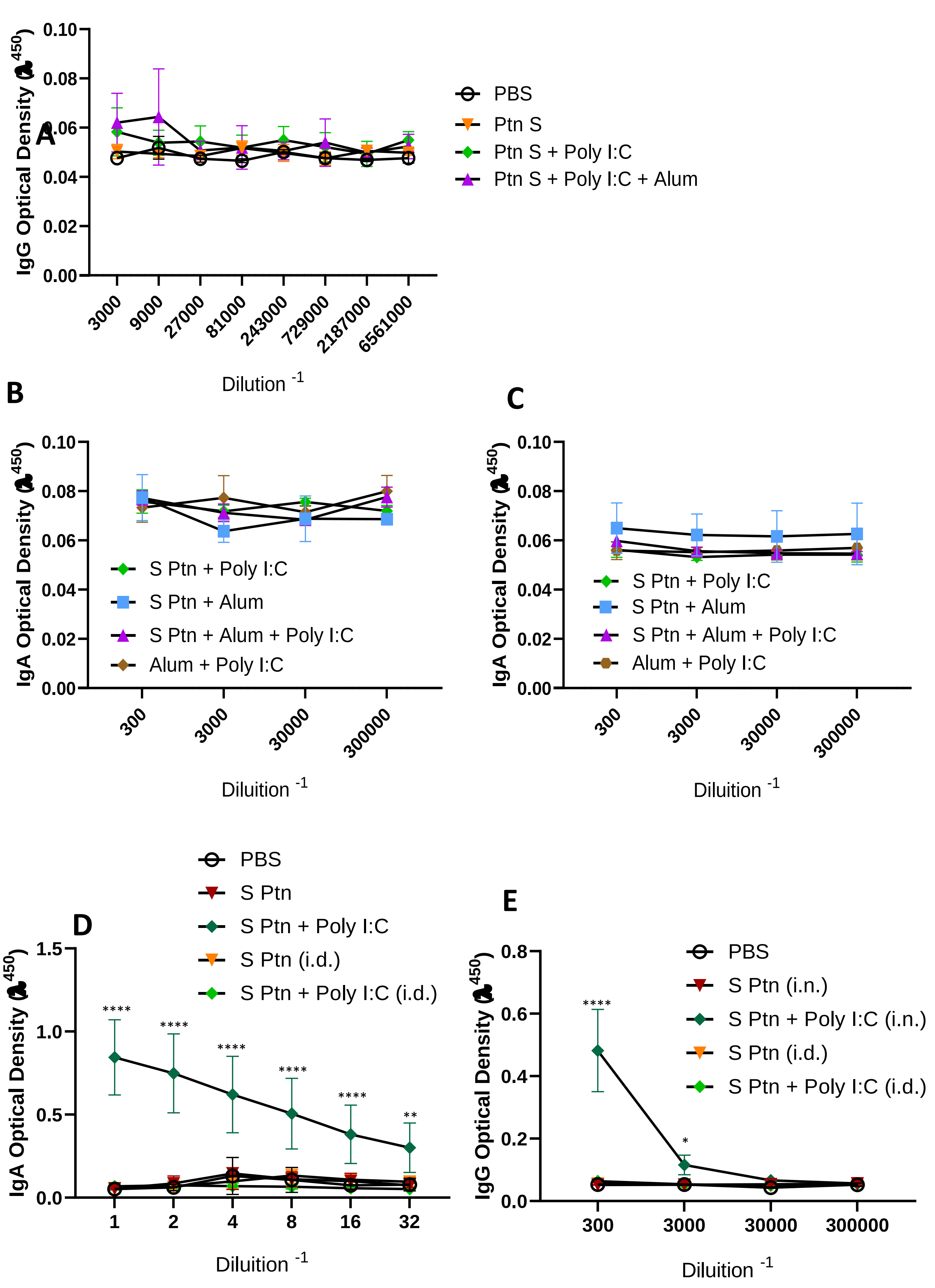

**Supplementary Figure 2**

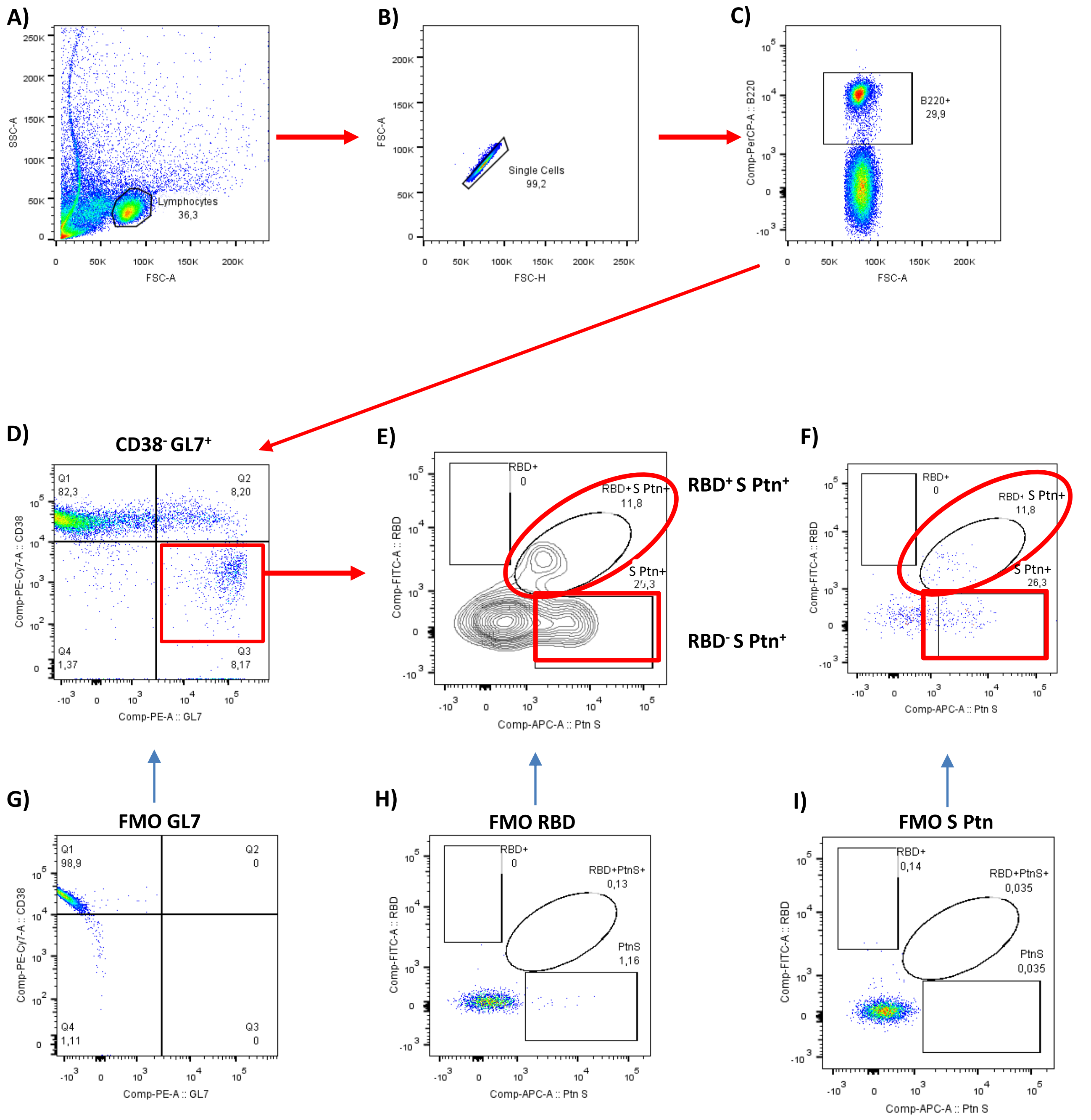

**Supplementary Figure 3**

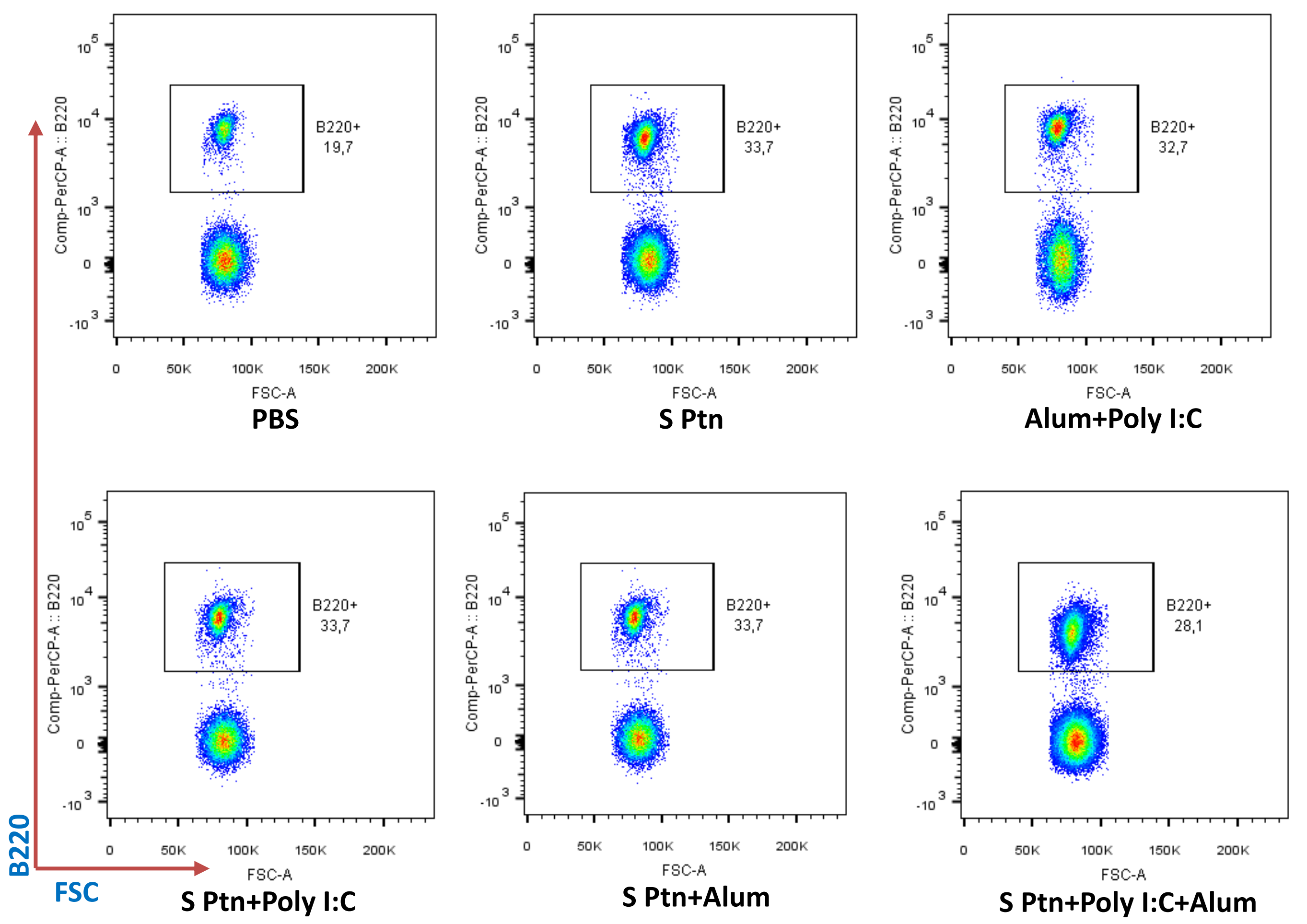

**Supplementary Figure 4**

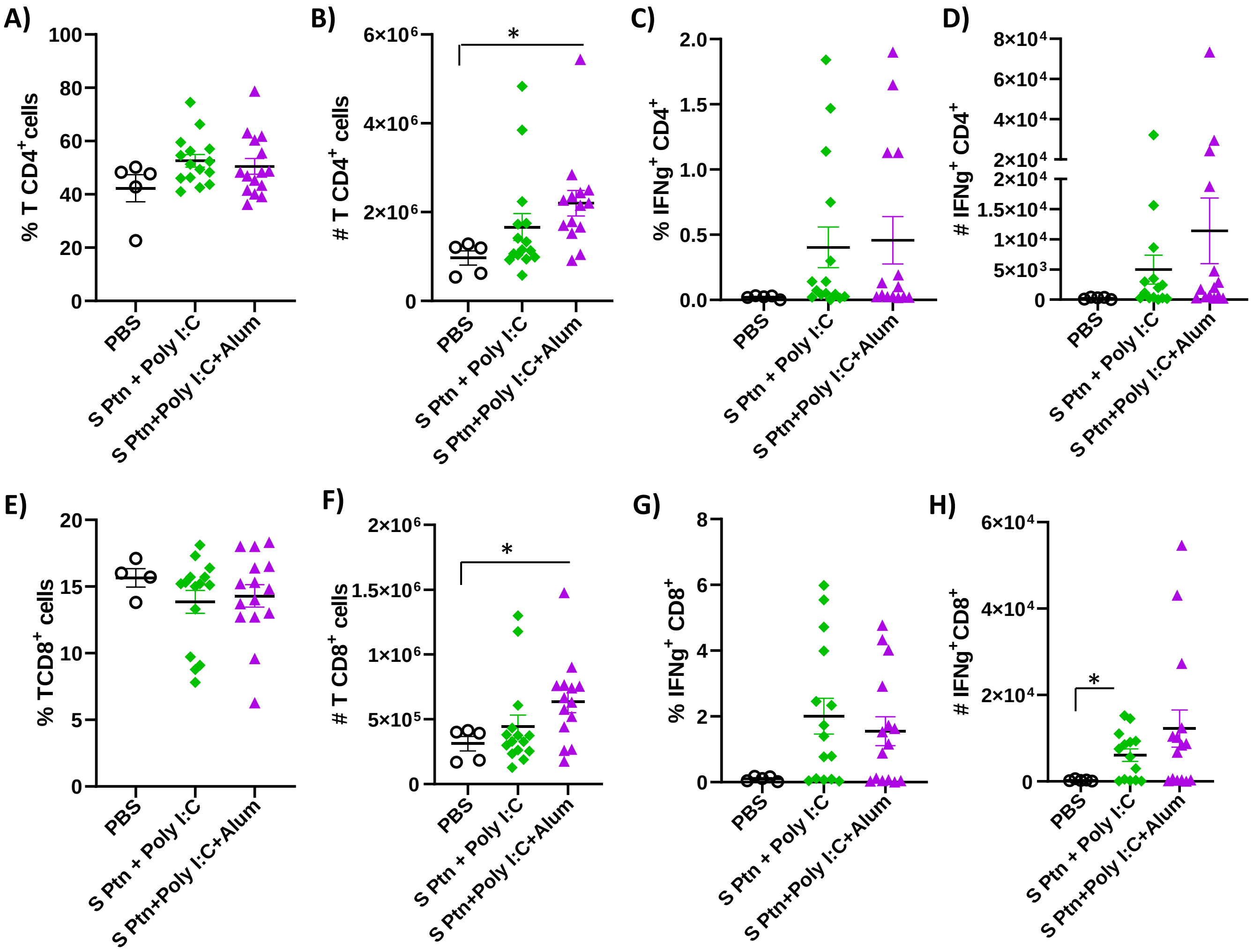

**Supplementary Figure 5**

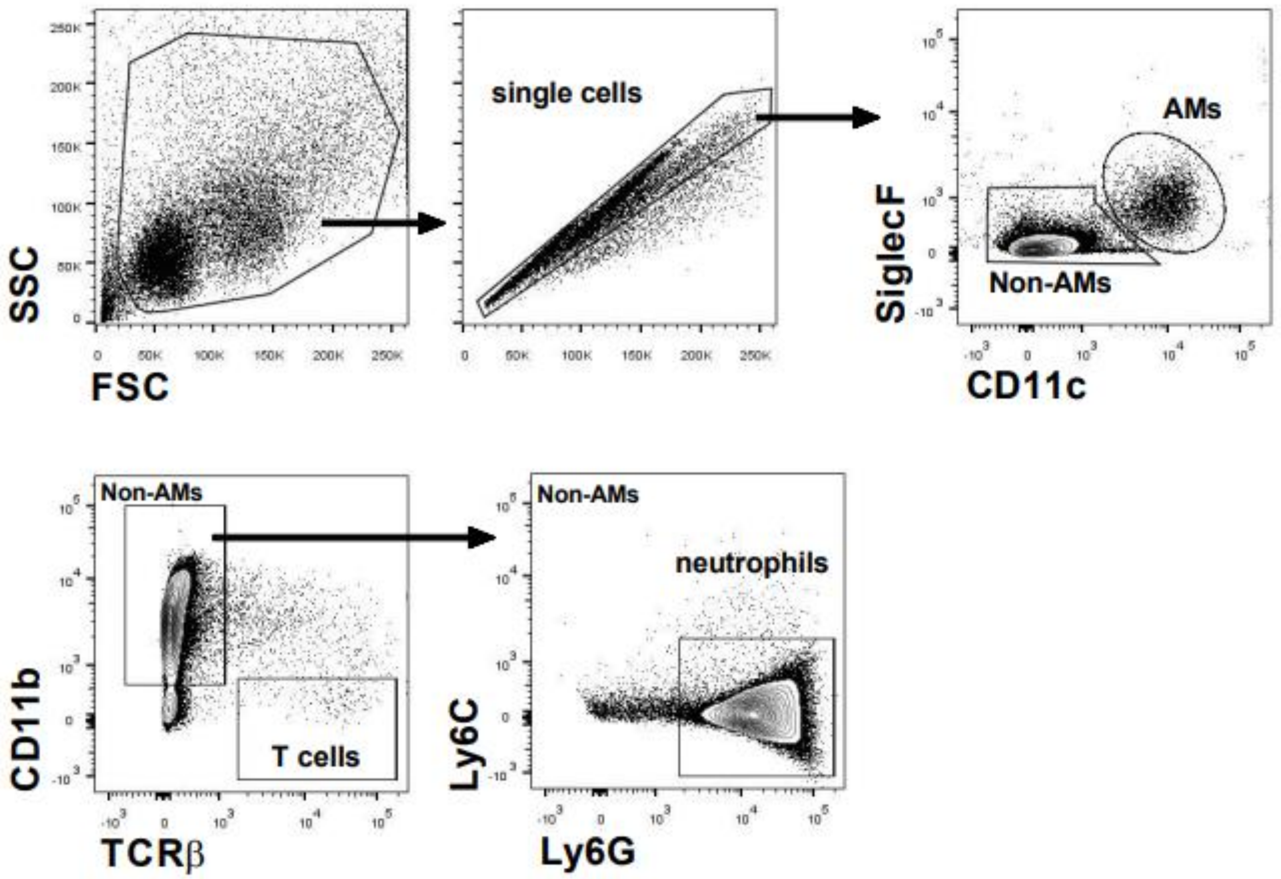

**Supplementary Figure 6**

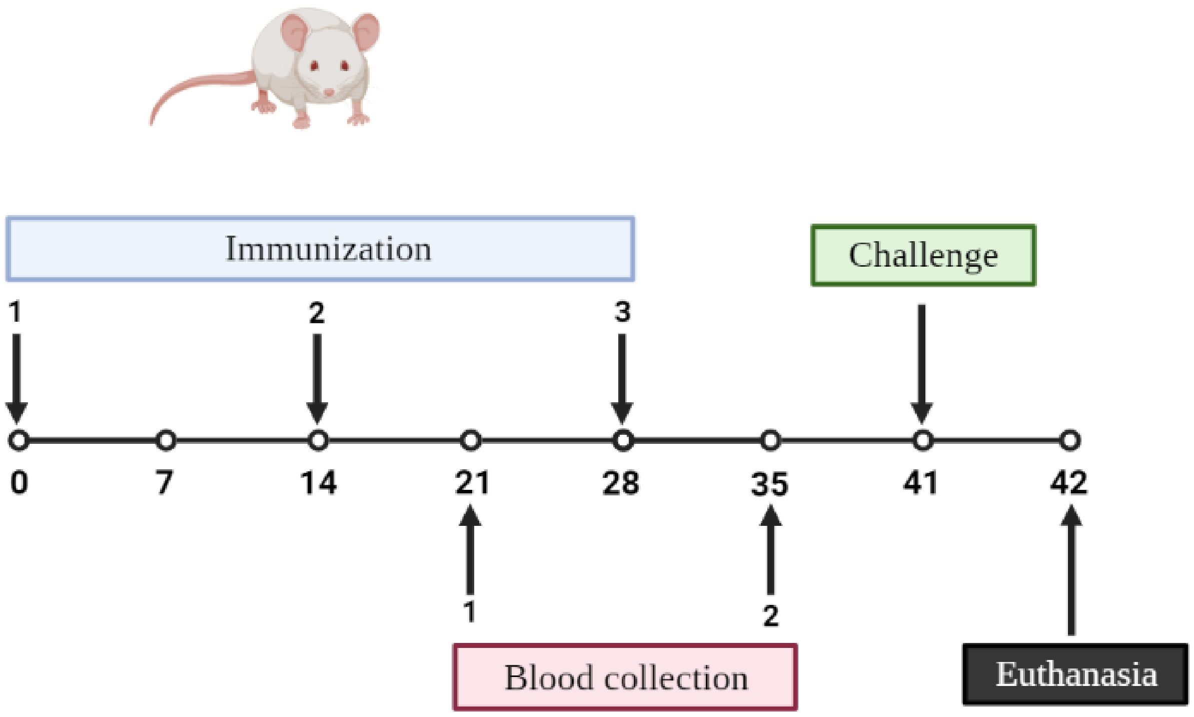
